# Endothelial CYB5R3 couples store-operated calcium entry to TRPV2 activation and vascular fitness

**DOI:** 10.1101/2025.10.08.681053

**Authors:** Mate Katona, Shuai Yuan, Robert Hall, Olivier Romito, Stefanie N. Taiclet, Sydney S. Tomman, Scott A. Hahn, Katherine Wood, Mohamed Trebak, Adam C. Straub

## Abstract

NADH–cytochrome b5 reductase 3 (CYB5R3) is a flavoprotein that governs nitric oxide (NO) signaling and supports NADPH oxidase 4–derived hydrogen peroxide production via coenzyme Q reduction in endothelium. While CYB5R3 expression is decreased during aging, the downstream consequences of CYB5R3 loss are not understood. Here, we demonstrate that depletion of CYB5R3 in primary human aortic endothelial cells activates a Ca^2+^ influx network characterized by the upregulation of calcium release-activated calcium (CRAC) channel subunits ORAI2 and ORAI3, as well as the non-selective cation channel transient receptor potential vanilloid 2 (TRPV2). When endoplasmic-reticulum Ca^2+^ stores were depleted, CYB5R3-deficient cells had increased Ca^2+^ entry through the plasma membrane, part of which was insensitive to classical store-operated Ca^2+^ entry (SOCE) blockers and was mediated by TRPV2, as demonstrated by genetic knockdown and pharmacologic inhibition. Mechanistically, loss of CYB5R3 increased Ca^2+^-dependent NO production through elevated CRAC channel activity, which oxidatively inhibited the protein tyrosine phosphatase non-receptor type 1 (PTPN1). This prevented TRPV2 dephosphorylation, thereby maintaining Janus kinase 1 (JAK1)-dependent channel activation downstream of SOCE. It also enhanced the responsiveness of TRPV2 to physiological heat stimuli. Thus, CYB5R3 normally acts as a brake, limiting NO-dependent PTPN1 oxidation and restraining TRPV2 activity. *In vivo*, endothelial-specific *Cyb5r3* deletion enhanced acetylcholine-induced vasorelaxation and improved exercise capacity, demonstrating a physiological function for this pathway in vascular adaptation. Together, these findings identify a CYB5R3–NO–SOCE– PTPN1–TRPV2 signaling axis that couples endothelial redox balance to Ca^2+^ dynamics and vascular function.

**SIGNIFICANCE:** Endothelial cells rely on receptor-regulated Ca^2+^ signals to produce vasodilators and control vascular function; however, the molecular mechanisms coordinating these pathways are incompletely understood. We identify CYB5R3 as a key redox switch that couples store-operated Ca^2+^ entry to the non-selective cation channel TRPV2. Loss of CYB5R3 enhances TRPV2 activity downstream of SOCE through NO-dependent oxidative inhibition of the phosphatase PTPN1, sustaining Janus kinase–mediated TRPV2 channel activation. This novel mechanism expands the physiological scope of CYB5R3 by redefining how redox enzymes intersect with Ca^2+^ signaling, linking endothelial CYB5R3 to vascular relaxation and exercise capacity *in vivo.* This positions CYB5R3 as a central regulator of vascular function with broad implications for cardiovascular health and disease.

## Methods

### Animals

All animal experiments were approved by and conducted in accordance with the University of Pittsburgh Institutional Animal Care and Use Committee. Cyb5r3 floxed mice (Cyb5r3^fl/fl^) were generated as previously described(1, 2). Both wild-type C57BL/6J and Cyb5r3^fl/fl^ mice were crossed with tamoxifen-inducible Cdh5(PAC)-CreERT2 mice (Figure 5A)(3). Male mice, aged 10–12 weeks, received tamoxifen (10 mg/kg) intraperitoneally for five days and were rested without treatment for seven days before the experiments. Mice were kept under a 12-hour light/dark cycle with free access to chow and water. All experiments were approved by the University of Pittsburgh Institutional Animal Care and Use Committee (IACUC), and all mice used are reported.

### Cell culture

Human aortic endothelial cells (HAEC) were purchased from Lonza and maintained in endothelial growth medium (EGM-2, Lonza, CC-3162) with 5% CO_2_ at 37°C. Following Lonza’s recommendation, population doubling was used to track the age of cells in culture, rather than the traditional method of using passage numbers. Cells with fewer than 13 population doublings were used for experiments.

### RNA sequencing

Human aortic endothelial cells (HAECs) were transfected with non-targeting and CYB5R3-targeting siRNA using Lipofectamine 3000 as described above. 48 hours after media change, cells were lysed for RNA extraction using the Qiagen RNeasy Mini kit. Total RNA samples were sent to Novogene for bulk RNA sequencing. Briefly, messenger RNA was purified from total RNA using poly-T oligo-attached magnetic beads for the preparation of an unstranded library. Libraries were pooled and sequenced on an Illumina platform to generate 150 bp paired-end reads. After removing adapter sequences, poly-N, and low-quality reads, clean reads were mapped to the reference genome using Hisat2 v2.0.5 and counted on the gene level using featureCounts v1.5.0-p3. Gene count data were filtered to retain genes with at least one count per million (CPM) in each sample. Differentially expressed genes (DEGs) between the two groups were determined with DESeq2 using adjusted p-value (false discovery rate [FDR]) < 0.05 as the threshold. DEGs were tested against the Gene Ontology Biological Process gene sets with clusterProfiler, and pathways with FDR < 0.05 were considered significantly overrepresented in DEGs. Selected significant pathways and related genes were used to generate a chord plot using the circlize package. The adjusted CPM (EdgeR) values were used to show the expression difference between the two groups.

### In vitro transfection of siRNA and expression vectors

The transfection of siRNA in HAECs was achieved using Lipofectamine 3000 (Thermo Fisher Scientific, L3000015) according to the manufacturer’s instructions. Non-targeting siRNA (siNT) and human Cyb5r3 targeting siRNA (siR3) were purchased from Horizon Discovery (D-001810-01-20 and L-009554-00-0005) and siNT and human TRPV2 from ThermoFisher. The day before transfection, HAECs were seeded at 15,000–20,000 cells/cm2 in 6-well plates. For Cyb5r3 or Trpv2 silencing alone, 10 μM of siNT or siR3 or siTRPV2 was given to HAECs in lipid complex overnight, before the medium was replenished to allow cells to recover. The incubation duration was counted from the time of medium change. For experiments involving Cyb5r3 and Trpv2 double knockdown, four groups of cells received 20 μM siNT, 10 μM siNT +10 μM siR3, 10 μM siNT +10 μM siTRPV2, or 10 μM siR3 + 10 μM siTRPV2, respectively. In this way, siRNA loads were kept the same in all treatment groups.

### Ca^2+^ Imaging

Human aortic endothelial cells treated with siRNA were seeded onto 25 mm Poly-D-lysine–coated coverslips (Sigma, P6407) at sub-confluence one day before imaging. Before the experiment, cells were incubated in extracellular medium (ECM; 140 mM NaCl, 4.9 mM KCl, 1.13 mM MgCl_2_, 10 mM HEPES, 10 mM D-glucose, 2 mM CaCl_2_, pH 7.4) and loaded with 2 µM Fura-2 AM (Invitrogen, F1221) for 20 minutes at room temperature. Epifluorescence Ca^2+^ imaging was performed using an Olympus IX51 inverted microscope equipped with a CoolLED pE-340fura LED illumination system (340/380 nm excitation), a dual-band dichroic and emission filter set (Chroma, 73100), and a Hamamatsu ORCA-Spark camera controlled by HCImage software. Imaging was conducted using a UV-optimized Olympus UAPO 40×/1.35 NA oil-immersion objective (UAPO/340). Intracellular Ca²⁺ levels were quantified as the ratio of Fura-2 fluorescence intensity at 340 nm and 380 nm excitation (F340/F380), representing relative changes in cytosolic Ca^2+^ concentration [Ca^2+^]_cyto_. Following dye loading, cells were washed and transferred into fresh ECM. SOCE protocol was performed as described in Figure 1B(1–4). Briefly, Fura-2-loaded cells were transferred to the stage in fresh ECM, then moved to 0 mM Ca^2+^ ECM 1 minute before imaging. After 1 minute of baseline recordings, cells were treated with 2 μM Thapsigargin (Tg) to empty the ER stores. After store depletion 5 mM Ca^2+^ was re-added to the ECM to trigger SOCE. After 3 minutes of SOCE induction, 5 μM Gd^3+^ was added to the ECM to block ORAI-dependent SOCE. At the end of the experiment, Fura-2 loading was tested by 10 μM Ionomycin. Traces from at least 3 independent experimental days were pooled and analyzed. Heat-induced Ca^2+^ transients were measured in Fura2-loaded HAECs. The chamber ECM was carefully replaced with pre-heated ECM to avoid shear stress-induced activation of ECs.

**Figure 1.**
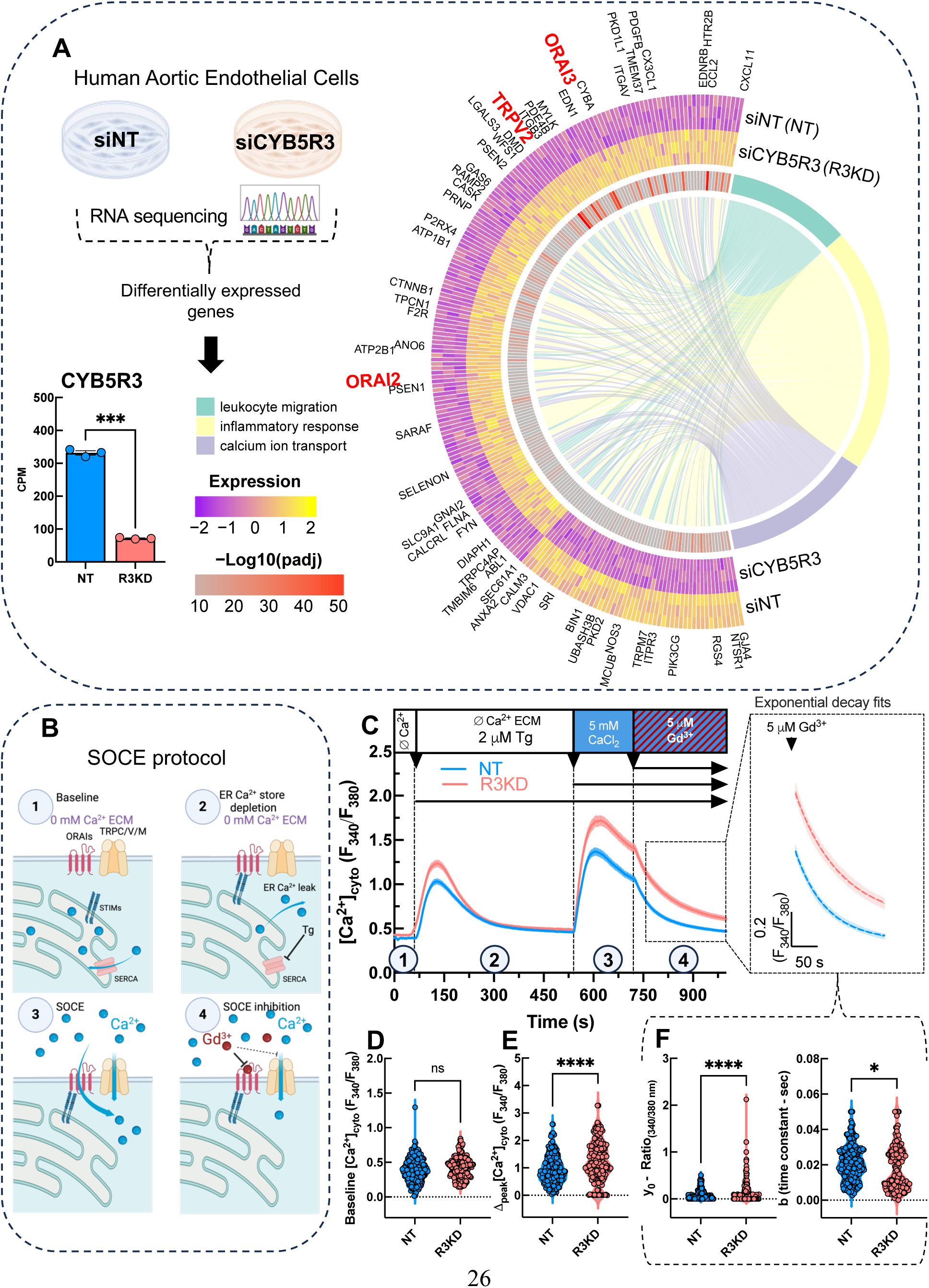
Loss of CYB5R3 altered endothelial Ca^2+^ signaling: A) RNAseq experimental design and siRNA knockdown efficacy (±SEM; n=3) with pathway analysis of differentially expressed genes in siNT (NT) and siCYB5R3-treated (R3KD) HAECs. B) Schematic describing the protocol used to study store-operated Ca^2+^ entry (SOCE): (1) bathing the cell in Ca^2+^-free extracellular media (ECM), (2) emptying the ER Ca^2+^ stores using Thapsigargin (Tg), (3) Ca^2+^ restoration to the bath solution to trigger SOCE, (4) blocking SOCE with 5µM Gd^3+^. C) Time courses (represented as mean [Ca^2+^]_cyto_ (F_340_/F_380_ Ratio) (± S.E.M.) showing the changes in [Ca^2+^]_cyto_ during the SOCE protocol in NT (n=230) control and R3KD (n=176) HAECs. D) Calculated baseline F_340_/F_380_ Ratio values of the individual traces. E) Individual Δpeak [Ca^2+^]_cyto_ values after Ca^2+^ re-addition to the ECM in NT and R3KD HAECs. F) Exponential decay fits showing the y0 and b (time constant) after Gd^3+^-induced inhibition of SOCE (calculated from the first 2 minutes of the inhibition). Statistics: Differential expression was analyzed using DESeq2 (Wald test with Benjamini–Hochberg FDR correction, FDR<0.05) ***p<0.0001. (A). Welch’s unpaired two-tailed t-test; NT vs R3KD, (D) ns (p=0.45), (E) ****p<0.0001, (F) y0 ****p<0.0001, b (Time Constant) *p=0.042.

### Whole cell patch clamp measurements:\

The whole-cell patch clamp protocol was optimized for HAECs, based on protocols previously described (4, 5). Briefly, HAECs were treated with siRNA as described above, then trypsinized and seeded on the bottom glass coverslips for 30 min, allowing cell adhesion. The media was washed using an extracellular solution containing 150 mM Na-gluconate, 10 mM glucose, 2 mM MgCl_2_, and 10 mM HEPES (pH adjusted to 7.4 with NaOH). Patch pipettes (2-5 MΩ) were pulled from borosilicate glass capillaries (World Precision Instruments) with a P-1000 Flaming/Brown micropipette puller (Sutter Instrument) and were filled with pipette solution containing 140 mM Cs-methanesulfonate, 2.5mM NaCl, 10mM EGTA, 10mM HEPES (pH adjusted to 7.2 using CsOH). We selected cells with small series resistance (< 10 MΩ) to perform the recordings. The cells were maintained at a holding potential of 0 mV before stimulation with a voltage ramp from –120 mV to +120 mV (lasting 800 ms, applied every 3 seconds). Recordings were performed with an Axopatch 200B and a Digidata 1440A (Molecular Devices) and were monitored with pCLAMP10. Data analysis was performed using Clampfit 10.1 software (https://www.moleculardevices.com/).

### *Ex vivo* wire myography

After tamoxifen treatments, mice were euthanized, and the aorta was collected for myography according to our previous publications (1). Briefly, thoracic aortae, mesentery and thoracodorsal arteries were rapidly excised placed in room temperature physiological salt solution (PSS), cleaned of fat, cut into 2 mm rings, and placed on a two-pin myograph (DMT 620M) filled with PSS containing (mM): NaCl 119, KCl 4.7, MgSO_4_ 1.17, KH_2_PO_4_ 1.18, D-glucose 5.5, NaHCO_3_ 25, EDTA 0.027, CaCl_2_ 2.5, pH 7.4 when bubbled with 95% O_2_ 5% CO_2_ at 37°C. Following a 30-minute rest, aortas were incrementally stretched to 500 mg initial tension. Vessels were then constricted with 60mM KCl for 5 minutes to test the viability and then washed three times with PSS and allowed to rest for 30 minutes. A final wash was performed, and vessels rested for an additional 10 minutes. Following the final 10-minute rest period, vessels were constricted with a dose-response of PGF2α (50uM-1uM). After reaching a plateau, a continuous dose-response curve of acetylcholine (10 nM-100 μM) was used to assess endothelium-dependent relaxation. Ca^2+^-free PSS containing 100μM sodium nitroprusside (SNP) was added to determine maximal dilation. The experimental protocol is outlined in Figure 5A.

### Immunoblotting

HAECs were lysed in radioimmunoprecipitation (RIPA) buffer with proteinase (P8340-5ML) and phosphatase inhibitors (P5726-5ML). Protein samples in RIPA buffer were quantified using Pierce™ BCA Protein Assay Kit (Thermo Fisher Scientific, 23225), diluted, and mixed with 5X Laemmli buffer. All samples were denatured at 95 °C for 10 minutes. Proteins were resolved in NuPAGE Bis-Tris gradient gels (Thermo Fisher Scientific, NP0336BOX) and transferred to 0.2 µm nitrocellulose membranes (BIO-RAD 1620112). Proteins of interest were probed with specific antibodies overnight at 4 °C, followed by infrared fluorescent secondary antibodies (IRDye® 680RD Donkey anti-Mouse IgG Secondary Antibody, LiCor #926-68072 or IRDye® 800CW Donkey anti-Rabbit IgG Secondary Antibody, LiCor #926-32213) for 90 minutes at room temperature. Blots were imaged with LI-COR Odyssey® CLx and BioRad Chemidoc^TM^, and fluorescent intensity was quantified with LI-COR Image Studio and BioRad ImageLab software. Antibodies and dilutions used in this study are listed in Table 1.

**Table 1.** List of primary antibodies used for immunoblotting. *Antibodies used for Ni-NTA pull-down.

| Epitope | Species | Dilution | Supplier | Catalog # |
| --- | --- | --- | --- | --- |
| Tubulin (alpha) | ms | 1:10000 | SigmaAldrich | T6074 |
| Jak1 | rb | 1:1000 | Cell Signalling | 3332S |
| Cytochrome B5 reductase 3 | rb | 1:1000 | Proteintech | 10894-1-AP |
| PTP1B | rb | 1:1000 | Abcam | AB244207-1001 |
| PTP1B (oxidized)* | ck | 2 µg/sample | MilliporeSigma | MABS456 |

### Ni-NTA Pull-down Experiments

To evaluate the oxidized fraction of PTPN1 in NT control and R3KD HAECs, we used an antibody (PTPN1-OX; Anti-PTP1B Antibody (Oxidized), Sigma-Aldrich) designed to detect the oxidation Cys215 oxidation-induced conformational changes of the PTPN1 catalytic domain. Cells were washed twice with cold (4°C) PBS and lysed in degassed lysis buffer (25 mM HEPES, pH 7.4, 150 mM NaCl, 0.25% deoxycholate, 1% TritonX-100, 25 mM NaF, 10 mM MgCl_2_, 1 mM EDTA, 10% glycerol, phosphatase (P5726, Roche) and protease inhibitor cocktail (P8340, Roche)). A subset of NT cells was incubated with H_2_O_2_ (100 μM) for 10 minutes, and a subset of both NT and R3KD cells was treated with L-NAME (100 μM) overnight before lysis. A subset of R3KD lysates was treated with DTT (1 mM) to ensure a post-lysis reducing environment. 1 mg lysate was treated with 2 µg of PTP1B-OX overnight at 4℃, gently rocking. Ni-NTA agarose (Qiagen, 50 μl as a 50% slurry equilibrated in binding buffer (20 mM HEPES, pH 7.4, 300 mM NaCl, 0.05% BSA, 0.05% Tween-20 and 10 mM imidazole) was added and incubated for one hour at 4°C. Protein complexes bound to Ni-NTA agarose beads were precipitated and washed (three times, 5 minutes each, at 4°C) with binding buffer containing 20 mM imidazole. The protein complexes were eluted from the Ni-NTA agarose beads with 500 mM imidazole (in binding buffer) for 15 minutes at 4°C with gentle shaking. Samples were run on SDS-PAGE, and total endogenous and oxidized PTPN1 were detected with an anti-PTPN1 antibody by immunoblotting.

### Quantification of gene expression

Cultured endothelial cells were directly lysed in the RLT buffer + β-Mercaptoetanol. Total RNA was extracted from RNeasy MiniElute Cleanup Kit (Qiagen, 74204) according to the manufacturer’s instructions. Reverse transcription was performed using SuperScript™ IV VILO™ Master Mix (Thermo Fisher Scientific, 11756050). Quantitative PCR for the cDNA library (RT-PCR) was set up with Power SYBR™ Green PCR Master Mix (Thermo Fisher Scientific, 4367659) and measured by QuantStudio^TM^ 5 real-time PCR machine (Thermo Fisher Scientific). The delta-delta Ct method was used to analyze mRNA expression levels. Primer sequences used in this study are listed in Table 2.

**Table 2.**
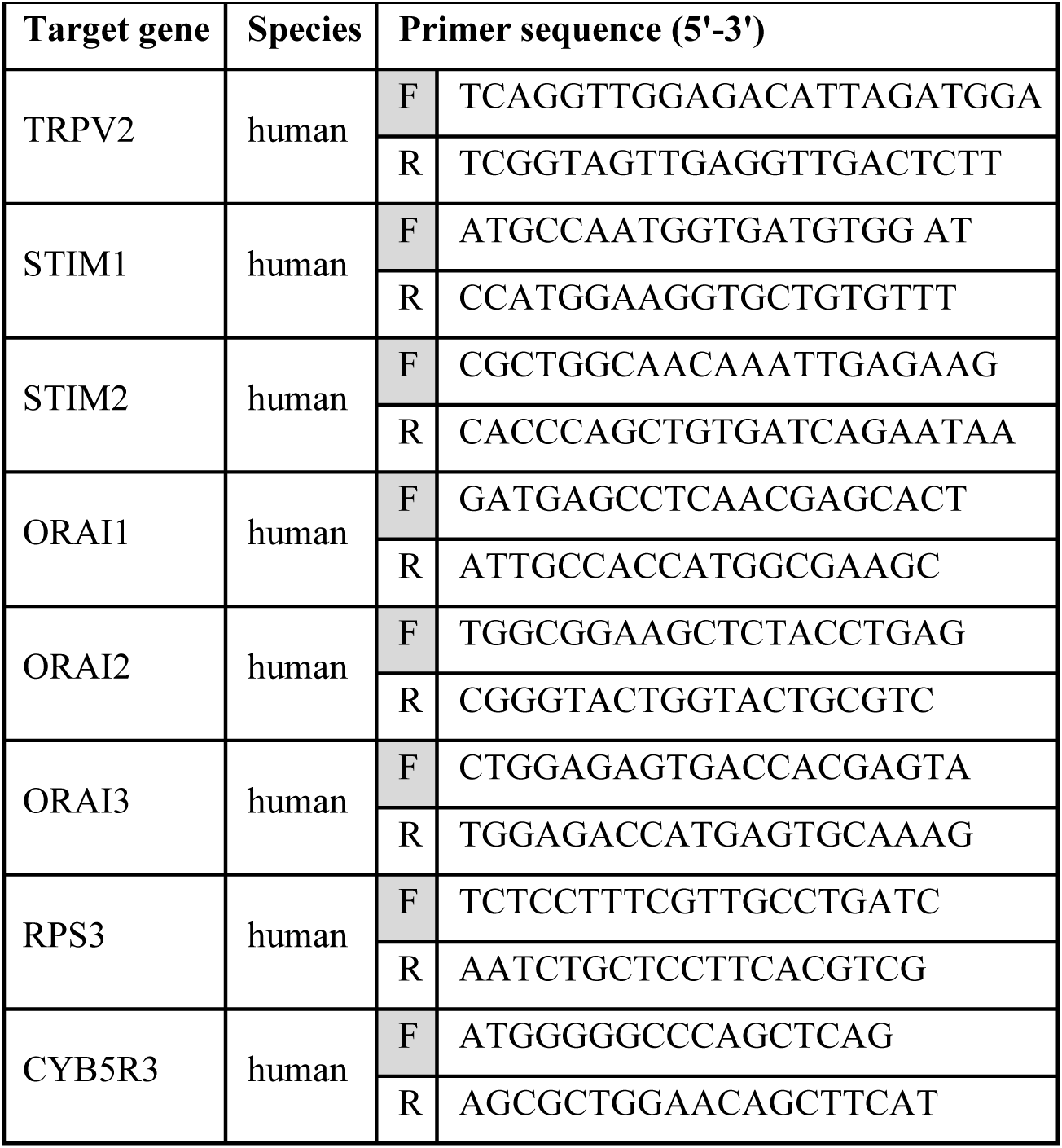
List of primer pairs used for Real-Time PCR.

### Exercise tolerance and blood flow measurements

Tamoxifen-injected mice were acclimated to treadmill exercise over a 5-day period by gradually increasing both the duration and speed of running sessions on an 8% incline. On the experimental day, animals were subjected to a treadmill endurance test (gradually increasing treadmill speed 0-18m/min over 30 minutes) and exercised until first exhaustion, defined as remaining in the fatigue zone (one mouse length at the bottom of the treadmill) for more than 5 seconds (**Figure 5E**). A separate cohort of animals was used for blood flow measurements using Laser Speckle Contrast Imaging (PeriCam PSI NR Laser Speckle Contrast Imaging, LSCI) of the lower limbs. After 5 days of treadmill acclimation, baseline LSCI measurements were obtained on day 1 on rested animals. The next day, the animals underwent a standardized 10-minute treadmill run (12m/min treadmill speed) after a 2-minute warmup, and post-exercise LSCI measurements were performed. For imaging, animals were anesthetized with isoflurane (3–4% for induction, 1–2% for maintenance in 100% oxygen) and maintained on a 37 °C heated pad. Imaging was performed right after anesthesia to reduce the vasodilatory effects of isoflurane. Regions of interest (ROIs) were set on the plantar surfaces of both paws. Measurements were taken at 25 frames/second and averaged every second until a stable perfusion rate was reached. Perfusion was averaged over the ROI within a two-minute measurement window. To further minimize the confounding effects of isoflurane on vascular reactivity, baseline and post-run measurements were taken on consecutive days.

## INTRODUCTION

NADH–cytochrome b5 reductase 3 (CYB5R3) is an evolutionarily conserved flavoprotein essential for cellular redox balance. Through electron transfer reactions, CYB5R3 supports heme reduction, lipid desaturation, cholesterol biosynthesis, xenobiotic metabolism, and the recycling of coenzyme Q (CoQ) (1, 2, 6–12). The membrane-bound isoform localizes to the endoplasmic reticulum, plasma membrane, and outer mitochondrial membrane, where it donates electrons either directly to substrates or indirectly through cytochrome b5 (7). Within the vasculature, CYB5R3 has emerged as a key regulator of nitric oxide (NO) signaling and redox homeostasis. It sustains NO bioactivity in resistance arteries by reducing endothelial α-globin heme iron, while in conduit arteries, it promotes NADPH oxidase 4-dependent hydrogen peroxide production through CoQ (2, 8).

Endothelial NO production is tightly controlled by cytosolic Ca^2+^ rise, primarily through Ca^2+^-calmodulin-dependent activation of endothelial nitric oxide synthase (eNOS) (13). Engagement of endothelial phospholipase C (PLC)-coupled receptors by growth factors and vasoactive agonists (e.g., angiotensin II, vasoactive intestinal peptide, acetylcholine) drives the hydrolysis of phosphatidylinositol-4,5-bisphosphate (PIP_2_) into inositol-1,4,5-trisphosphate (IP_3_) and diacylglycerol (DAG). IP_3_ triggers Ca^2+^ release from the endoplasmic reticulum (ER), and depletion of ER stores activates ER-resident stromal interaction molecules (STIM), which aggregate in ER-plasma membrane junctions to trap and gate ORAI hexameric channels and trigger store-operated Ca^2+^ entry (SOCE) (14–18). The biophysical manifestation of SOCE is the highly Ca^2+^ selective current termed Ca^2+^ release-activated Ca^2+^ (CRAC) (19, 20). Furthermore, non-selective transient receptor potential (TRP) channels are also activated downstream of PLC signaling by various means, including cytosolic Ca^2+^ and lipid second messenger, and often amplify cytosolic Ca^2+^ elevations initiated by SOCE. CRAC channels not only replenish Ca^2+^ stores but also promote transcriptional, metabolic, and inflammatory vascular and airway remodeling (15, 21–23). In contrast, vascular TRP channels integrate diverse stimuli, including mechanical, thermal, and oxidative inputs (24). For instance, the Ca^2+^-activated Na^+^-conducting TRP melastatin 4 (TRPM4) channels in smooth muscle regulate myogenic tone by causing depolarization and subsequent activation of Ca_V_1.2 channels (25, 26). In the brain, endothelial TRP ankyrin 1 (TRPA1) is activated by 4-hydroxy-nonenal to augment Ca^2+^ influx and cause vasorelaxation through the activation of IK_Ca_ and SK_Ca_ potassium channels (27, 28). TRP polycystin (TRPP) channels in vascular smooth muscle and endothelia are activated by intravascular pressure and flow, respectively, to mediate vasoconstriction and vasodilation (29–32). While these pathways highlight the versatility of Ca^2+^ regulation in the vasculature, the roles of other TRP channels, which are abundant in the endothelium, remain incompletely defined.

To investigate the functional consequences of decreased CYB5R3 expression during aging, we conducted transcriptomic profiling of primary human aortic endothelial cells (HAECs) with CYB5R3 knockdown. Results revealed that CYB5R3 knockdown alters the expression of Ca^2+^ transport–related genes, including ORAI2, ORAI3, and the non-selective cation channel TRPV2, as well as Janus kinase 1 (JAK1) and protein tyrosine phosphatase non-receptor type 1 (PTPN1), two regulators of TRPV2 activity (33). Functional studies demonstrated that CYB5R3 loss triggers a TRPV2-dependent Ca^2+^ influx pathway downstream of CRAC channel activation. Mechanistically, this pathway is sustained by NO-dependent oxidative inhibition of PTPN1, which prevents TRPV2 dephosphorylation and permits JAK1-mediated TRPV2 channel activation. Collectively, these findings identify CYB5R3 as a redox regulator of endothelial Ca^2+^ signaling, uncovering a previously unrecognized mechanism by which the endothelium adapts to mechanical and thermal stress to preserve vascular function.

## RESULTS

### CYB5R3 regulates aortic endothelial Ca^2+^ signaling

CYB5R3 is broadly expressed across the vasculature; however, endothelial α-globin, a primary target of CYB5R3, is primarily confined to resistance arteries (8). To elucidate the broader redox-regulatory role of CYB5R3 beyond its canonical functions in α-globin (8) and CoQ (2) reduction, we performed RNA sequencing in HAECs, which lack α-globin expression. This approach enabled us to assess the effects of siCYB5R3 knockdown (R3KD) on endothelial gene expression independent of α-globin.

Our analysis revealed the differential expression of 1,768 genes resulting from the knockdown of CYB5R3. Pathway analysis revealed that several genes associated with Ca^2+^ homeostasis were impacted, including channels involved in transporting Ca^2+^ across the plasma membrane into the cytosol **(Figure 1A)**. Two members of the CRAC channels family, ORAI2 and ORAI3, key players in SOCE, and the transient receptor potential type vanilloid 2 (TRPV2), which has an elusive role in vascular endothelial cells, were upregulated. We used epifluorescence microscopy to measure changes in cytosolic Ca^2+^ concentration ([Ca^2+^]_cyto_) to determine whether the transcriptional changes related to R3KD influenced Ca^2+^ homeostasis in HAECs. Cells were loaded with the Ca^2+^ dye Fura2 and treated in the absence of extracellular Ca^2+^ with thapsigargin (Tg), an inhibitor of the sarco/endoplasmic reticulum Ca^2+^-ATPase (SERCA), which passively depletes ER Ca^2+^ stores. This was followed by restoration of 5 mM Ca^2+^ to the extracellular milieu to record the magnitude of SOCE **(Figure 1B&C; 1-3)**. We found no differences in the baseline of Fura2 F_340_/F_380_ ratios, suggesting no differences in resting [Ca^2+^]_cyto_ between siRNA non-targeting (NT) controls and R3KD **(Figure 1D)**. However, SOCE was significantly elevated in R3KD cells compared to NT controls **(Figure 1E)**. The contribution of ORAIs to this SOCE was examined using the relatively low concentration (5 µM) of Gd^3+^, a specific blocker of CRAC channels **(Figure 1B-C;4)**. While Tg-induced Ca^2+^ influx was elevated in R3KD cells compared to the NT control cells, this influx was only partially inhibited by 5 µM Gd^3+^ **(Figure 1C;4)** and by the pharmacological SOCE inhibitor, GSK7975A at 10 µM **(Supplementary Figure 1A-C)**. An exponential decay fit on the Gd^3+^ and GSK7975-induced inhibition confirmed this partial inhibition, suggesting that an additional Ca^2+^-conducting channel might not be affected or only partially blocked by these SOCE inhibitors **(Figure 1F)**. We then tested whether the contribution from this additional Ca^2+^-conducting channel to thapsigargin-activated Ca^2+^ influx required the activation of CRAC channels. Therefore, we pre-incubated cells with 5 µM Gd^3+^ to completely block CRAC channels prior to initiation of recordings. This protocol prevented thapsigargin-activated Ca^2+^ influx in Ca^2+^-containing extracellular bath solutions. This suggests that CRAC channel activation is a prerequisite for the activation of this additional Ca^2+^ influx mechanism in the R3KD cells **(Supplementary Figure 1D-E)**.

### TRPV2 is activated downstream of CRAC channel activity

TRPV2, a plasma membrane Ca^2+^ channel, was upregulated at the transcript level in R3KD HAECs **(Figure 1A)**. Thus, we investigated the potential contribution of TRPV2 to the additional Ca^2+^ influx pathway triggered downstream SOCE in R3KD cells. We knocked down TRPV2 (NT&TRPV2KD) either alone or in conjunction with R3KD (R3&TRPV2). Knockdown of TRPV2 restored the Gd^3+^-sensitivity to the Ca^2+^ influx phase across all conditions, suggesting that this Ca^2+^ influx is now purely mediated by SOCE **(Figure 2A&B)**. Like most non-selective TRP channels, TRPV2 itself was also reported to be sensitive to trivalent lanthanides, such as Gd^3+^ and La^3+^, however, at a substantially higher concentration of 100 µM (34). Interestingly, both TRPV2KD and the double R3&TRPV2 KD conditions showed enhanced SOCE **(Figure 2A)**. We used quantitative reverse transcription PCR (qPCR) to determine the expression of SOCE proteins in TRPV2-deficient cells. We found significant upregulation of stromal interaction molecules 1 and 2 (STIM1 and 2) and ORAI2 transcripts **(Supplementary Figure 2A)**, indicating that increased SOCE in TRPV2KD cells resulted from the upregulation of STIM and ORAI gene expression and that this SOCE enhancement is independent of TRPV2 function **(Figure 2A)**. Furthermore, acute TRPV2 inhibition by tranilast (TRN) also restored the Gd^3+^ sensitivity of thapsigargin-activated Ca^2+^ influx, further supporting that TRPV2 is the contributor to the additional Ca^2+^ influx downstream of SOCE **(Supplementary Fig. 2B&C)**. The acute pharmacological inhibition of TRPV2 did not affect SOCE **(Supplementary Fig. 2D)**, suggesting that the enhanced SOCE upon TRPV2KD is due to long-term transcriptional upregulation of STIM and ORAI genes **(Supplementary Fig. 2A)**, independently of the loss of CYB5R3.

**Figure 2.**
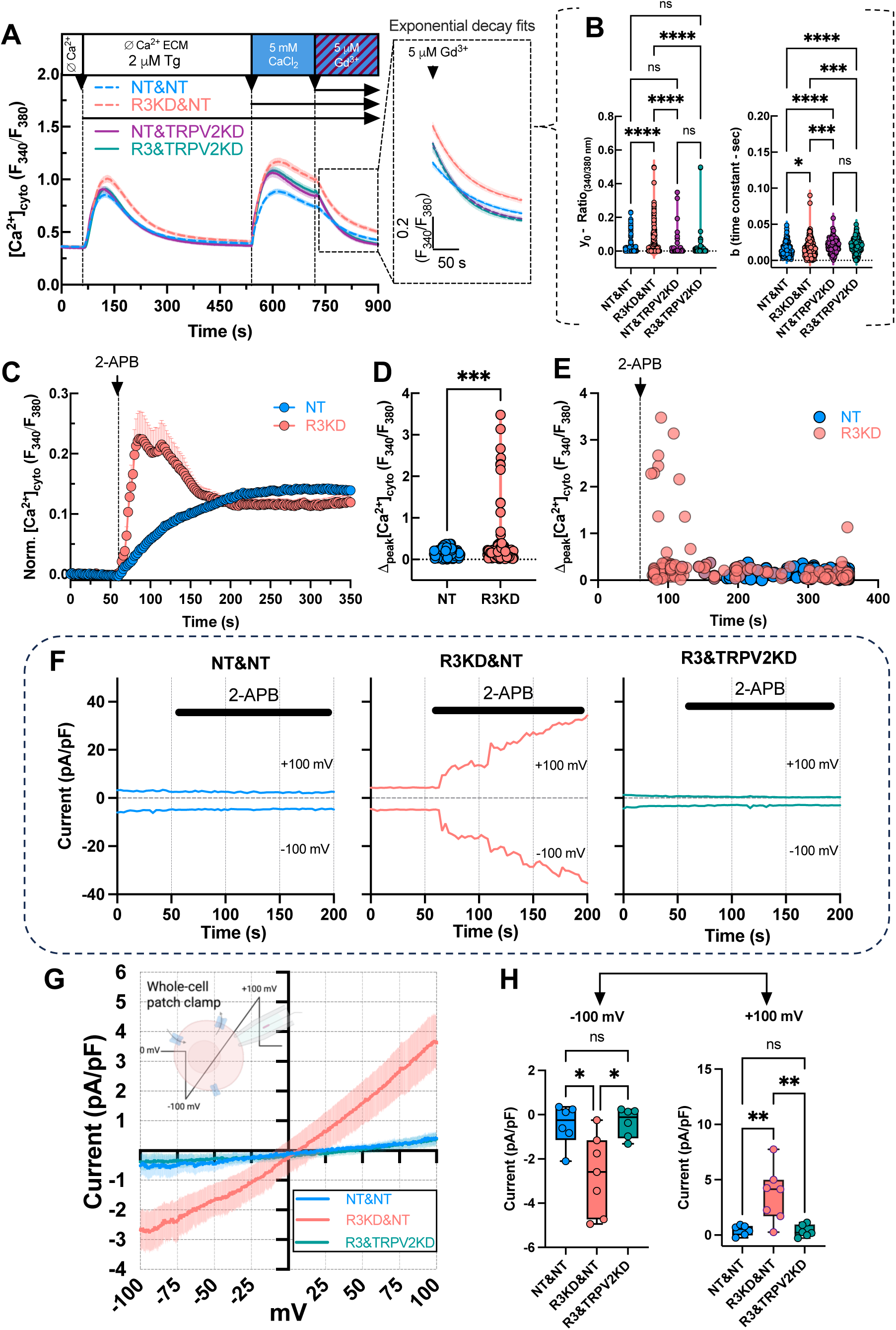
2-APB-induced cationic currents in R3KD cells that were lost on additional TRPV2 KD and TRPV2 KD restored the Gd^3+^ sensitivity of SOCE in R3KD cells: A) Mean (± S.E.M.) time traces showing the changes in [Ca^2+^]_cyto_ (F_340_/F_380_ Ratio) during SOCE protocol in double siRNA treated siNT_Dharmacon_ and siNT_Thermo_ (NT&NT) control siNT_Thermo_ and siCYB5R3 treated (R3KD&NT), siNT_Dharmacon_ and siTRPV2 treated TRPV2KD (NT&TRPV2KD) and siCYB5R3 and siTRPV2KD (R3&TRPV2KD) HAECs. B) Exponential decay fits showing the y0 and b (time constant) after Gd^3+^-induced inhibition of SOCE (calculated from the first 2 minutes of the inhibition). C) Normalized mean time courses (± S.E.M.) of the 300 μM 2-APB-induced changes of [Ca^2+^]_cyto_ in NT and R3KD HAECs. D) Δ[Ca^2+^]_cyto_ readings represented as violin plots with all data points visualized. E) Δ[Ca^2+^]_cyto_ readings plotted at the time of the peak to show the time distribution. F) Representative time courses of 300 μM 2-APB-induced cation currents (at +100 mV and −100 mV) in whole-cell mode in NT&NT, R3KD&NT, and R3&TRPV2KD HAECs (Y scales matched in all graphs). G) Average I/V relationships in NT&NT, R3KD&NT, and R3&TRPV2KD HAECs in response to 300 μM 2-APB. H) Steady state cation current density (at +120 mV and −120 mV) represented as boxplots with the min to max with all data points, mean, and median for each condition. **Statistics:** (B) One-way ANOVA with Šídák’s multiple comparisons test; (y0) NT&NT vs. R3KD&NT, R3KD&NT vs. NT&TRPV2KD, R3KD&NT vs. R3&TRPV2KD ****p<0.0001, others ns p>0.9999, (b) NT&NT vs. R3KD&NT *p=0.0306, R3KD&NT vs. NT&TRPV2KD, **p=0.0027, R3KD&NT vs. R3&TRPV2KD, ***p=0.0008, NT&NT vs. NT&TRPV2KD, NT&TRPV2KD vs. R3&TRPV2KD, ****p<0.0001, NT&TRPV2KD vs. R3&TRPV2KD ns p>0.9999. D) NT:0.160, *n*=167; R3KD:0.323, *n*=129; unpaired two-tailed t-test *t*(294)=3.45, *\*\*\*p*=0.0006; mean difference: 0.163 ± 0.047, 95% CI: 0.070–0.256, η²=0.039). H) Whole-cell currents at –100 mV and +100 mV. At –100 mV, one-way ANOVA revealed significant differences among groups (F(2,15) = 5.625, *P* = 0.015, R² = 0.43) with no evidence of unequal variances (Brown–Forsythe *P* = 0.051; Bartlett’s *P* = 0.091). Tukey’s test showed that R3KD&NT currents differed significantly from both NT&NT (*\*p*=0.041) and R3&TRPV2KD (*\*p*=0.020), whereas NT&NT and R3&TRPV2KD were not different (*P*=0.93). At +100 mV, ANOVA also detected significant differences (F(2,15)=10.39, *P*=0.0015, R²=0.58), although variance tests indicated unequal SDs (Brown–Forsythe *P*=0.038; Bartlett’s *P*=0.0004). Tukey’s test revealed that R3KD&NT currents were significantly different from NT&NT (*\*\*p*=0.0031) and R3&TRPV2KD (*\*\*p*=0.0039), whereas NT&NT and R3&TRPV2KD did not differ (*P*=0.99).

To directly test plasma membrane TRPV2 activity, we stimulated cells with 300 μM 2-aminoethoxydiphenylborate (2-APB), a known activator of TRPV2 channels, and recorded Ca^2+^ transients (**Figure 2C–E**) and whole-cell cationic currents (**Figure 2F–G**). Control double siNT (NT&NT)-treated cells showed a slow elevation in the [Ca^2+^]_cyto_ and negligible cation currents. In contrast, a fraction of R3KD cells (52.7% vs 0% NT) exhibited robust 2-APB–induced fast ([Ca^2+^]_cyto_ peak <60s after 2-APB addition) Ca^2+^ transients and TRPV2-like non-selective currents (+3.62 ± 2.47 S.D. pA/pF at +100 mV; –2.7 ± 1.78 S.D. pA/pF at –100 mV; R3KD n=7/13 cells vs +0.43 ± 2.0.46 S.D. pA/pF at +100 mV; –0.47 ± 0.93 pA/pF at –100 mV; NT n=2/12 cells) with a reversal potential near 0 mV. Importantly, these currents were abolished in R3&TRPV2 double KD cells (+0.37 ± 0.54 S.D. pA/pF at +100 mV; –0.36 ± 0.66 S.D. pA/pF at –100 mV; R3&TRPV2KD n=0/12 cells), suggesting that TRPV2 mediates these currents. Although the densities of these TRPV2 currents were modest, they represent native currents recorded from primary human endothelial cells, underscoring the physiological relevance of R3KD-enhanced TRPV2 activity and Ca^2+^ signaling in endothelial cells.

### R3KD led to the emergence of thermo-sensitive Ca^2+^ signals in HAECs

TRPV2 orthologs in rodents are activated by temperature, but this property is considered less prominent in human TRPV2 (35–37). However, most studies investigating the thermal regulation of human TRPV2 have been conducted in overexpression models using cell types that primarily do not express endogenous TRPV2 (37). Consequently, potential accessory molecules and subcellular organizations could have been missing, which may be essential for the thermo-sensitivity of the human TRPV2 ortholog. We showed that CRAC channel activation increased TRPV2 activity in R3KD HAECs **(Figure 2A)**. We aimed to determine whether R3KD influenced human endothelial heat-activated Ca^2+^ responses, a known activator of TRPV channels (33–35). Unlike control cells, a large proportion of R3KD cells (26.6% R3KD vs 4.6% NT) exhibited rapid cytosolic Ca^2+^ transients when challenged by a temperature of ∼ 42℃ **(Figure 3A&G)**. This oscillatory response was absent in TRPV2 KD conditions. A larger proportion (40%) of the cells in TRPV2 KD conditions exhibited a delayed response during the cooling phase (**Figure 3B&G**). This response likely reflects the previously described heat-off activation of SOCE (38), which is upregulated in TRPV2 KD cells **(Supplementary Figure 2A)**. These data indicate that the presence of TRPV2 is necessary to generate the early heat-activated fast transients **(Figure 3B&G)**. Mo et al. demonstrated on murine macrophages that JAK1 and PTPN1 regulated the heat sensitivity threshold of TRPV2 by mediating the phosphorylation of distinct tyrosine motifs (33). These motifs are conserved in both murine and human TRPV2 orthologs **(Supplementary Figure 3A)**. Interestingly, we observed the upregulation of JAK1 transcripts **(Supplementary Figure 3B)** and proteins **(Supplementary Figure 3C&D)** and the downregulation of PTPN1 transcripts **(Supplementary Figure 3B)** in R3KD cells. Therefore, we tested whether increased tyrosine phosphorylation could modify the thermal sensitivity of TRPV2. We treated control and R3KD cells with Upadacitinib (UPA), a specific inhibitor of JAK1 activity **(Figure 3C.a.)**(39–41). We found that 6 hours of UPA treatment completely abolished the heat-induced Ca^2+^ transients in R3KD cells **(Figure 3D&G)**. While the resting [Ca^2+^]_cyto_ was not affected by either siRNA treatment or pharmacological inhibitors **(Figure 3E)**, the peak [Ca^2+^]_cyto_ was elevated in the R3KD cells **(Figure 3F)**. We then examined whether JAK1 inhibition affected TRPV2 activation downstream of SOCE activation by thapsigargin **(Figure 3C.b.)**. UPA treatment decreased thapsigargin-activated Ca^2+^ influx **(Figure 3H&J)** while restoring its Gd^3+^ sensitivity **(Figure 3I**), suggesting that in the presence of the JAK1 inhibitor UPA, this thapsigargin-activated Ca^2+^ influx is now purely mediated by SOCE. These findings indicate that human TRPV2 channels are regulated by heat and tyrosine phosphorylation in a way similar to its mouse ortholog.

**Figure 3.**
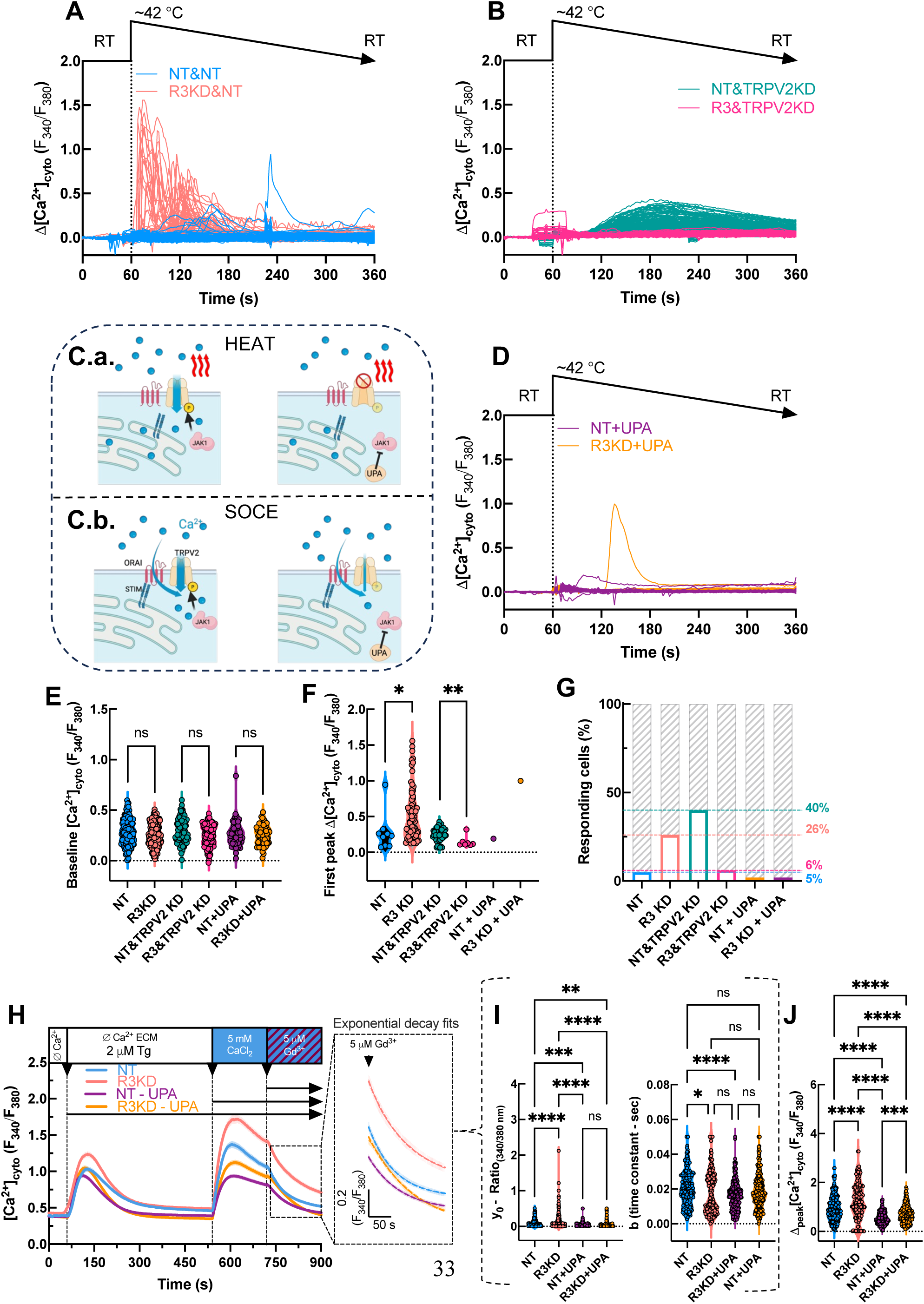
Emergence of heat-activated Ca^2+^ signals in R3KD cells, which were lost upon TRPV2 KD or JAK1 pharmacological inhibition: (A) Individual time courses of Δ[Ca^2+^]_cyto_ (F_340_/F_380_ Ratio) changes induced by ∼42 ℃ extracellular medium (ECM) in double siRNA-treated siNT and siNT (NT&NT) control siNT and siCYB5R3-treated (R3KD&NT) and (B) siNT and siTRPV2 treated TRPV2KD (NT&TRPV2KD) and siCYB5R3 and siTRPV2KD (R3&TRPV2KD) HAECs. (C) Schematic showing the proposed mechanism of TRPV2 post-translational modifications caused by JAK1 and the effect of JAK1 inhibition. (D) Individual time courses of Δ[Ca^2+^]_cyto_ (F_340_/F_380_ Ratio) changes induced by ∼42℃ECM in upadacitinib (UPA) treated NT control and R3KD cells. (E) Individual baseline [Ca^2+^]_cyto_ (F_340_/F_380_ Ratio) values. (F) Calculated Δ[Ca^2+^]_cyto_ (F_340_/F_380_ Ratio) peak values of the first heat-induced Ca^2+^ responses. (G) The percentage (%) of cells responding to heat. (H) Mean time traces showing the changes in [Ca^2+^]_cyto_ (F_340_/F_380_ Ratio) during SOCE protocol in siNT control and siCYB5R3 treated HAECs with and without UPA treatment. (I) Exponential decay fits showing the y0 and b (time constant) after Gd^3+^-induced inhibition of SOCE(calculated from the first 2 minutes of the inhibition). (J) Individual Δ[Ca^2+^]_cyto_ (F_340_/F_380_ Ratio) peak values after Ca^2+^ re-addition. **Statistics:** (E) Welch’s unpaired two-tailed t-test; NT vs R3KD, NT&TRPV2 KD vs R3&TRPV2 KD, NT+UPA vs R3KD + UPA, all comparisons ns. (F) Welch’s unpaired two-tailed t-test; NT vs R3KD *p=0.0216, NT&TRPV2 KD vs R3&TRPV2 KD **p=0.0124. (I) One-way ANOVA, F(5, 1012) = 23.80, ***P < 0.0001; variances unequal. Šídák’s: R3KD > all others (***P < 0.0001)

### Increased SOCE in R3KD HAECs induced eNOS phosphorylation

Endothelial Ca^2+^ signaling increased NO production through enhanced eNOS activity. By adapting the SOCE protocol from Figure 1C(1–3), we studied the effects of increased SOCE on eNOS(S1177) phosphorylation (p-eNOS) and downstream NO generation using immunoblotting in R3KD cells after 5 minutes of extracellular Ca^2+^ restoration. We found increased p-eNOS/total-eNOS ratios in R3KD cells **(Figure 4A&B)**, indicating enhanced Ca^2+^-induced eNOS activity and NO production. Pre-blocking SOCE with 5 µM Gd^3+^ prevented the SOCE-dependent eNOS phosphorylation. These findings demonstrate that SOCE controls NO production and that loss of CYB5R3 amplifies Ca^2+^-induced eNOS activation and NO production in endothelial cells, highlighting a critical role for CYB5R3 in regulating endothelial Ca^2+^ signaling and redox balance—processes essential for maintaining vascular tone and function.

**Figure 4.**
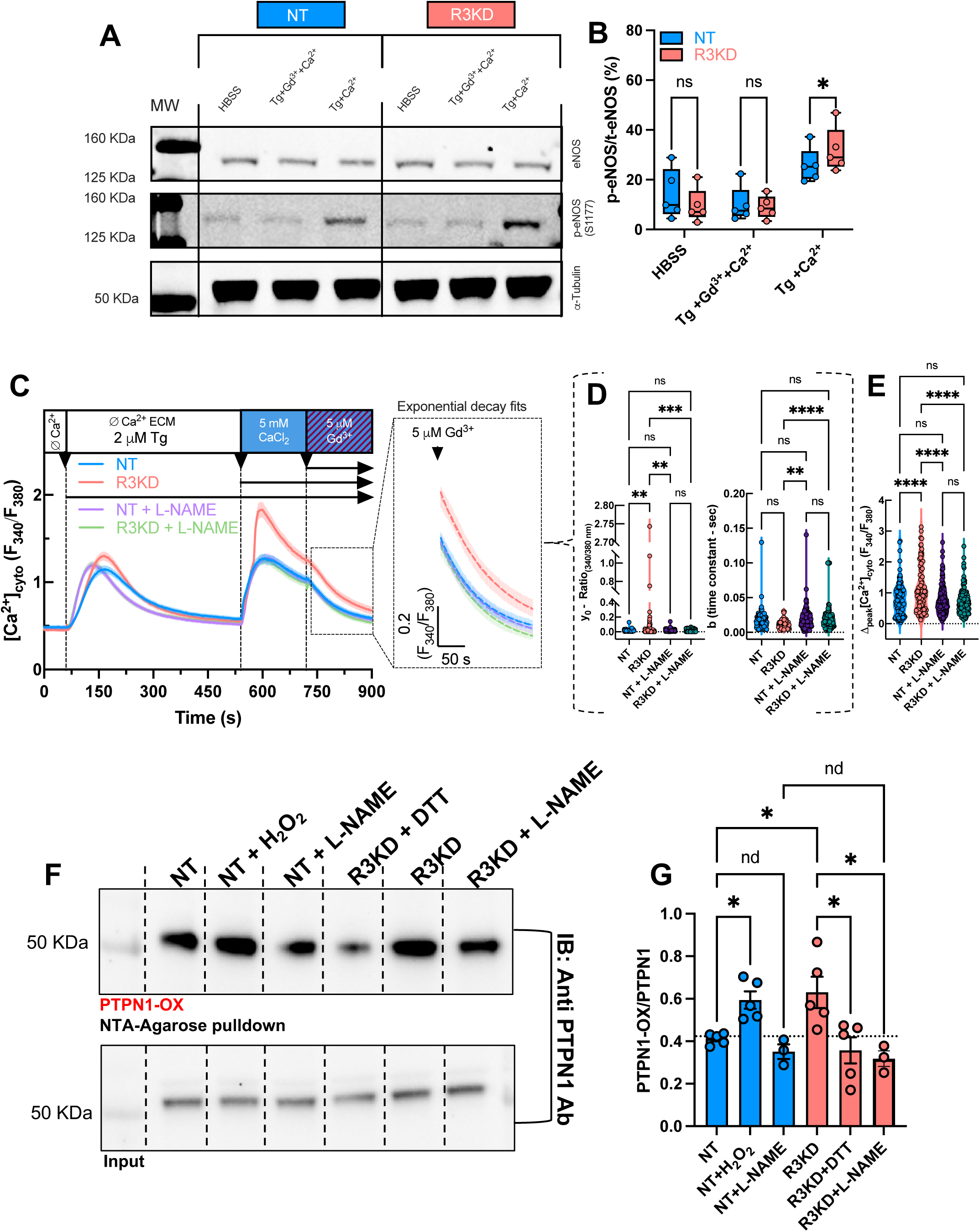
NO-dependent inactivation of PTPN1 enhances TRPV2 activity in R3KD cells: (A) Representative immunoblots and (B) quantification of p-eNOS(S1177)/total eNOS in NT and R3KD HAECs ± Gd^3+^ (5 µM) (n=5. (C) [Ca^2+^]_cyto_ traces in NT and R3KD cells ± L-NAME (100 µM), showing restored Gd^3+^ sensitivity in R3KD. (D) Exponential decay fits Y0 and b (Time Constants). (E) Δ[Ca^2+^]_cyto_ (F_340_/F_380_ Ratio) peak values after Ca^2+^ re-addition. (F) Representative Ni-NTA pulldown of oxidized PTPN1 (Cys215-OX) in NT and R3KD cells ± DTT or L-NAME. (G) quantification of PTPN1-OX/PTPN1. **Statistics:** (A) Unpaired t-test, t(8) = 5.05, *P = 0.0010; Δ mean = 22.91 ± 4.53. (B) One-way ANOVA, F(3, 489) = 5.62, P = 0.0009; Šídák’s: R3KD > NT, NT+L-NAME (**P < 0.01), R3KD+L-NAME (***P = 0.0008); others ns. (D) y0: F(3,489)=5.62, p=0.0009; NT vs R3KD **p=0.0086, R3KD vs NT+L-NAME **p=0.0038, R3KD vs R3KD+L-NAME ***p=0.0008. b (Time Constant): F(3,489)=6.75, *p*=0.0002; R3KD vs NT+L-NAME \*\**p*=0.0067, R3KD vs R3KD+L-NAME \*\*\*\**p*<0.0001; all other comparisons ns. (E) One-way ANOVA, F(3,489)=13.52, p<0.0001; NT vs R3KD ***p<0.0001, R3KD vs NT+L-NAME ***p<0.0001, R3KD vs R3KD+L-NAME ***p<0.0001; all other comparisons ns. (G) Kruskal–Wallis P = 0.0028; R3KD > NT (*P = 0.026), reversed by DTT or L-NAME (*P < 0.01); H_2_O_2_ mimicked R3KD, rest of the comparisons showed no difference (nd).

**Figure 5.**
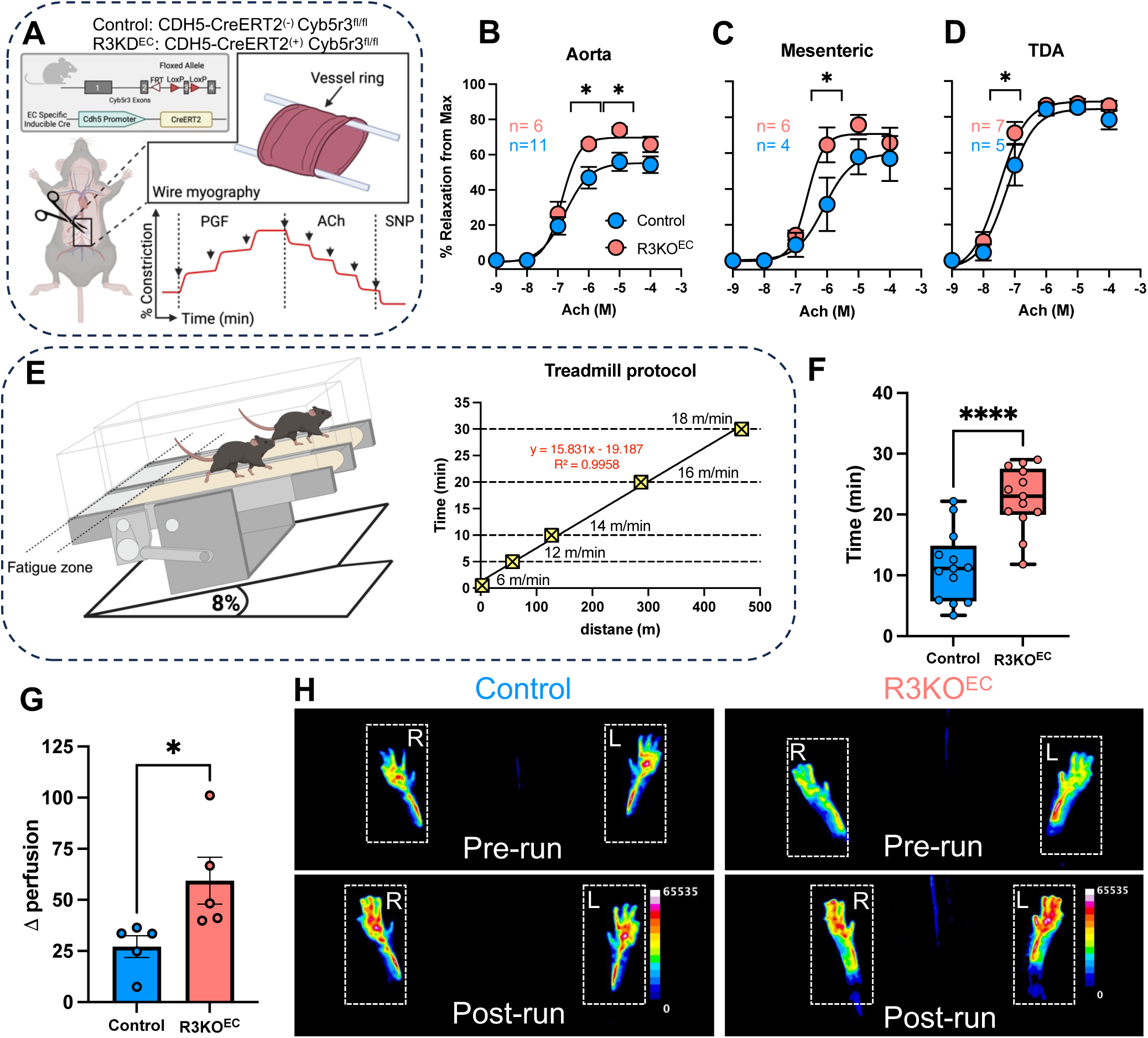
Endothelial-specific Cyb5r3 KO mice have increased ACh-induced vasorelaxation, exercise tolerance, and exercise-induced blood flow: (A) Animal model and myography protocol schematic. (B-D) ACh-induced vasorelaxation in the aorta, mesenteric artery, and TDA, shown as % relaxation. (E) Schematic of the Treadmill fatigue test with increasing speed and 8% incline. (F) Exercise tolerance time. (G) Δ perfusion (Post–Pre run). **Statistics:** (B-D) Two-way ANOVA: genotype effects in (B) aorta (P = 0.0010), (C) mesenteric (P = 0.0131), and TDA (P = 0.028); ACh dose ***P < 0.0001; no interaction (P > 0.16). Šídák’s post hoc: KO > control at higher doses (P < 0.05); others ns. E) Unpaired t-test, t(24) = 5.19, **P < 0.0001; Δ = 11.24 ± 2.16. G) Unpaired t-test, t(8) = 2.56, P = 0.034; Δ = 32.27 ± 12.63.

### Inhibition of endothelial NO normalized Ca^2+^ influx in R3KD HAECs

To determine whether eNOS activity and NO production play a role in Ca^2+^ signaling of R3KD HAECs, we treated the cells with 100 µM L-NAME for 6 hours and then measured changes in [Ca^2+^]_cyto_ using the thapsigargin protocol **(Figure 4C-E)**. Our findings showed that L-NAME treatment completely normalized the enhanced SOCE of R3KD cells, and restored the inhibition of Ca^2+^ influx by 5 µM Gd^3+^ **(Figure 4D)**, suggesting that this Ca^2+^ influx is now purely mediated by SOCE and that L-NAME likely restricts TRPV2 activation downstream of SOCE **(Figure 4E)**. Importantly, L-NAME treatment did not affect NT control cells, further supporting the idea that only the portion of thapsigargin-activated Ca^2+^ influx that is mediated by TRPV2 in R3KD cells is NO-dependent.

### Loss of CYB5R3 decreased PTPN1 activity through NO-dependent thiol modification

NO is primarily recognized as a signaling molecule with antioxidant and vasodilatory properties. However, it can also act as a pro-oxidant under certain conditions by interacting with reactive oxygen species (ROS) and reactive nitrogen species (RNS). Our group previously demonstrated that the loss of CYB5R3 resulted in elevated superoxide (O_2-_) levels in HAECs due to the lack of NOX4-derived O_2-_ conversion to H_2_O_2_ at the mitochondria (2). These alterations in the redox environment of HAECs may affect the function of redox-sensitive proteins. Our RNAseq dataset indicated transcriptional changes in JAK1 and PTPN1 **(Supplementary Fig. 3B)**. While JAK1 phosphorylates TRPV2 to enhance its activity, PTPN1 has been shown to dephosphorylate TRPV2 and JAK1(33), thereby reducing their activity. Endothelial NO and O_2_^−^ can interact to form peroxynitrite, a potent inactivator (among other pro-oxidants) of PTPN1 (42, 43). Some of these redox modifications have been reported to be irreversible; however, these studies were performed on isolated enzymes with non-physiological concentrations of pro-oxidants. We studied oxidized PTPN1 (PTPN1-OX)(44) at Cys215 using nickel-nitrilotriacetic acid (Ni-NTA) pulldown assays. R3KD cells exhibited elevated levels of oxidized PTPN1, indicating that the altered redox environment inactivated PTPN1 **(Figure 4F&G)**, potentially leading to increased TRPV2 tyrosine phosphorylation and enhanced activity. To test the reversibility of oxidized PTPN1 in R3KD cells, we treated the lysates with 1 mM DTT and observed a decrease in the oxidized PTPN1 band. This suggests that PTPN1 Cys215 is likely modified by S-nitrosation. To further clarify the role of increased endothelial NO production in this process, we treated the cells with 100 µM L-NAME overnight to block eNOS activity. We found lower amounts of oxidized PTPN1, confirming that PTPN1 oxidation in R3KD HAECs was NO-dependent. These data suggest that CYB5R3 disrupts redox homeostasis in HAECs, leading to increased NO-dependent reversible oxidation of PTPN1 at Cys215, which inactivates its phosphatase activity, enhances TRPV2 phosphorylation, and increases channel activity to either heat or downstream of SOCE activation—effects that are reversible by reducing agents and are dependent on eNOS-derived NO.

### Ablation of endothelial Cyb5r3 in mice increased fitness and vasodilatation in various vascular beds

To study the effects of endothelial Cyb5r3 KO (R3KO^EC^) on acetylcholine-dependent vasorelaxation, we performed ex vivo myography on vessels from the aorta, mesenteric, and thoracodorsal beds of Cdh5(PAC)-Cre(+), Cyb5r3^fl/fl^ KO and Cdh5(PAC)-Cre(–), Cyb5r3^fl/fl^ control mice **(Figure 5A)**. After pre-constriction with prostaglandin F2⍺ (PGF; 10^−8^-10^−5^ M), vessels received increasing concentrations of acetylcholine (ACh; 10^−9^–10^−4^ M). ACh activates muscarinic receptors, raising intracellular Ca^2+^ and stimulating eNOS to produce NO and EDHF for vasorelaxation. Arteries from R3KO^EC^ mice showed greater ACh-induced vasorelaxation than controls, indicating that CYB5R3-deficient endothelial cells respond more effectively to ACh by increasing Ca^2+^ and NO production for vasorelaxation in various vascular beds. **(Figure 5B-D)**. To test the physiological consequence of increased Ca^2+^-dependent vasorelaxation in R3KO^EC^ animals, we pre-trained littermate control and KO animals on a treadmill for 5 days before conducting the exhaustion experiment. Muscle work generates excess heat and increases blood flow to support the higher O_2_ demand. These are all physiological triggers of endothelial Ca^2+^signaling and subsequent vasorelaxation. On the day of the experiment, we ran the animals at increasing treadmill speeds until they first exhibited exhaustion, as determined by spending over 5 seconds in the ‘fatigue zone’ at the bottom of the treadmill, located above the electric stimulus grid **(Figure 5E)**. We evaluated the total time spent on the treadmill before reaching exhaustion and found that the R3KO^EC^ animals spent nearly twice the time on the treadmill compared to their littermate controls. To test whether this difference was due to elevated blood flow from increased vasorelaxation, we measured the blood flow of R3KO^EC^ and littermate controls using laser speckle contrast imaging (LSCI). The animals were pre-trained to use the treadmill and run simultaneously on the experimental day. We measured baseline blood flow the day before to reduce the vasoactive effect of isoflurane used for anesthesia during LSCI. We calculated the exercise-induced changes in the blood flow of the animals. We found that the R3KO^EC^ animals had elevated blood flow in their lower limbs after exercise compared to Control animals **(Figure 5G&H)**. These results are consistent with our hypothesis that Cyb5r3 loss triggers a Ca^2+^/NO positive feedback loop involving TRPV2 channels that enhances NO production and subsequent vasorelaxation, increasing blood flow to supply the higher oxygen demand of the working muscles, ultimately resulting in increased performance.

## DISCUSSION

The regulation of vascular tone requires precise integration of redox homeostasis and Ca²⁺ signaling, orchestrated by redox-sensitive switches that govern vasodilator synthesis (45–47). Building on prior work, here we identify CYB5R3 as a redox checkpoint that suppresses endothelial Ca²⁺ influx through both SOCE and TRPV2 activity, acting via an NO-phosphatase feedback loop. This work expands CYB5R3’s role beyond coenzyme Q recycling (2) and α-globin heme reduction (8), placing it at the nexus of Ca^2+^ influx, phosphatase regulation, and vascular adaptation.

### CYB5R3 Links SOCE and TRPV2

Transcriptomic profiling of CYB5R3 knockdown endothelial cells revealed enrichment of Ca^2+^ transport pathways **(Figure 1A)**. Functionally, CYB5R3 deficiency augmented thapsigargin-activated Ca^2+^ influx, making it partially resistant to CRAC channel inhibitors **(Figure 1B–F; Supplementary Fig. 1D&E)**, which implicates an additional Ca^2+^ entry pathway activated downstream of SOCE. Silencing and pharmacologic inhibition of TRPV2 non-selective cation channel restored the Gd^3+^ sensitivity of thapsigargin-activated Ca^2+^ influx, implicating TRPV2 in mediating this additional Ca^2+^ influx pathway **(Figure 2A–C; Supplementary Fig. 2B-D)**. Interestingly, TRPV2 knockdown also upregulated STIM1/2 and ORAI2 **(Supplementary Fig. 2A)**, suggesting compensatory transcriptional crosstalk that maintains Ca²⁺ entry. Acute TRPV2 inhibition with tranilast also restored the Gd^3+^ sensitivity of thapsigargin-activated Ca^2+^ influx without altering SOCE amplitude **(Supplementary Fig. 2B–D)**.

We further demonstrate that 2-APB activates endogenous human TRPV2, eliciting rapid Ca²⁺ transients **(Figure 2C–E)** and whole-cell currents in CYB5R3-deficient cells **(Figure 2F–H)**. Although 2-APB affects ORAI3 (48) and other TRPV channels (49), siRNA knockdown confirmed that TRPV2 is the mediator of 2-APB-activated Ca^2+^ transients and membrane currents in CYB5R3-deficient endothelial cells.

Other TRPs modulate CRAC activity (50) TRPV1-derived Ca^2+^ microdomains can inactivate ORAI1 (51), TRPV4 associates with STIM1/ORAI1 to shape flow-induced oscillations (52, 53), and TRPV6 influences ORAI1/STIM1 trafficking (54). Our findings extend this paradigm by suggesting TRPV2 amplifies CRAC channel function, with CYB5R3 serving as a molecular brake to maintain endothelial Ca^2+^ balance.

### Thermal and Phosphorylation-Dependent Gating of TRPV2

A conceptual advance of this work is the demonstration that native human endothelial TRPV2 is heat-sensitive when appropriately tuned by phosphorylation of conserved tyrosine motifs **(Supplementary Figure 3A)** (33). Although human TRPV2 has been viewed as functionally inactive in response to heat in overexpression models (36, 37, 55), we found that CYB5R3 loss induced oscillatory Ca^2+^ transients when cells were challenged by ∼42 °C **(Figure 3A),** which were abolished by TRPV2 knockdown **(Figure 3B)** or modulators of tyrosine phosphorylation **(Figure 3D)**. Mechanistically, CYB5R3 silencing increased JAK1 and downregulated PTPN1 transcription **(Supplementary Fig. 3B&C)**, favoring a phosphorylated TRPV2 state (33). JAK1 inhibition abolished both the Gd^3+^-insensitive portion of thapsigargin-activated Ca^2+^ influx **(Figure 3H-J)** and the thermal activation of Ca^2+^ transients, consistent with the characteristics of murine TRPV2 channels (33, 56) and demonstrating that JAK1 may serve as a dynamic switch for TRPV2 gating in endothelial cells.

### A Ca^2+^–NO–TRPV2 Feedback Loop

Enhanced Ca^2+^ influx in CYB5R3-deficient cells increased eNOS phosphorylation and NO production **(Figure 4A&B)**. Inhibition of eNOS normalized the magnitude of SOCE in CYB5R3-deficient endothelial cells and restored its full Gd^3+^ sensitivity **(Figure 4C&D)**, indicating that NO sustains TRPV2 hyperactivity. This loop converges on PTPN1 (42–44, 57), a redox-sensitive phosphatase regulating JAK (58) and TRP channel phosphorylation (33, 59–61). CYB5R3 loss promoted the reversible oxidation of PTPN1 at Cys215, impairing its catalytic function and enabling persistent TRPV2 phosphorylation **(Figure 4F&G)**. These redox modifications were reversed by reducing agents and prevented by NO inhibition, indicating that NO-derived oxidants drive PTPN1 inactivation.

### *In Vivo* Consequences of *Cyb5r3* Deletion

Endothelial-specific *Cyb5r3* deletion enhanced acetylcholine-induced vasorelaxation across aortic, mesenteric, and thoracodorsal arteries **(Figure 5B-D)**. This phenotype translated into increased treadmill endurance **(Figure 5F)** and enhanced exercise-induced lower limb perfusion **(Figure 5G)**, demonstrating improved NO-dependent vasodilation and enhanced functional vascular reserve. Reduced CYB5R3 expression, observed during aging (11) and in patients with hypomorphic variants (62, 63), may thus serve as an early compensatory mechanism to preserve perfusion under stress. However, chronic TRPV2 hyperactivation could lead to maladaptive remodeling or abnormal angiogenesis. Interpretation is further complicated by ⍺-globin, which limits NO bioavailability through heme iron-dependent scavenging (64, 65), and by EDHF-mediated pathways in resistance arteries (45). Enhanced relaxation observed in Cyb5r3-deficient vessels reflects integrated NO and EDHF contributions, rather than singular CYB5R3–TRPV2 coupling effect. Moreover, NO itself can upregulate TRPV2 expression in the brain (66, 67), raising the possibility that increased NO bioavailability secondarily enhances TRPV2 activity, further amplifying the vascular phenotype.

### Broader Physiological and Therapeutic Implications

Beyond the vasculature, TRPV2 is expressed in the brain, immune cells, cardiomyocytes, and tumors, where it influences cell survival, stress responses, and proliferation. Our findings identify phosphorylation and redox sensitivity as generalizable mechanisms of TRPV2 gating, potentially explaining how this channel integrates oxidants (66–68), mechanical stress (69, 70), cannabinoids (56, 71), and temperature (33, 36, 37). Systemic cardiovascular effects associated with TRPV2 ligands including altered blood pressure and tachycardia (72, 73) may therefore reflect activation of the CYB5R3–NO–PTPN1 pathway. The CYB5R3 T117S (c.350C>G) variant, common in individuals of African ancestry (∼23% allele frequency, rare in other populations), encodes a partial loss-of-function enzyme with ∼50–60% activity and has been linked to increased cardiovascular risk (62, 74–76). Because endothelial CYB5R3 loss enhances TRPV2 activity, carriers of T117S may similarly exhibit augmented TRPV2-mediated Ca^2+^ influx and altered redox–Ca^2+^ coupling that extend beyond endothelial cells to other cell types, including arterial and cardiac myocytes. Notably, tranilast, a TRPV2 inhibitor, has been shown to improve or stabilize cardiac function and reduce TRPV2 expression in muscular dystrophy–associated heart failure (77, 78), underscoring the therapeutic potential of targeting this pathway in muscle cells. Although direct evidence linking T117S to TRPV2 hyperactivation and Ca^2+^ dysregulation in cardiac and skeletal muscle has yet to be established, these findings collectively suggest a plausible mechanistic connection that warrants further study.

### Conclusion

We identify CYB5R3 as a redox gatekeeper that integrates Ca^2+^ entry, phosphatase activity, and NO signaling to maintain endothelial balance. Three conceptual advances emerge: (i) CYB5R3 regulates SOCE and TRPV2-mediated Ca^2+^ influx; (ii) CYB5R3 prevents NO-driven oxidation of PTPN1, thereby restraining TRPV2 hyperactivation; and (iii) endothelial CYB5R3 loss enhances vasodilation and exercise performance *in vivo* **(Figure 6)**. By positioning TRPV2 within this redox-sensitive axis, our study extends its physiological relevance beyond thermal sensing and highlights it as a potential therapeutic target. While CYB5R3 loss in endothelial cells enhances vascular fitness, the loss of CYB5R3 in other vascular cell types, including vascular smooth muscle and cardiomyocytes might have opposite effects, warranting thorough investigations. Restoring CYB5R3 activity or directly modulating TRPV2 phosphorylation could rebalance Ca^2+^-NO signaling in cardiovascular disease. Given that CYB5R3 expression declines with aging and cardiovascular disease, dysregulation of this axis may contribute to endothelial dysfunction. Depending on the disease, therapeutic strategies that either enhance or inhibit CYB5R3 activity or TRPV2 channel function may offer new opportunities to correct Ca^2+^-NO imbalance in vascular disease.

**Figure 6:**
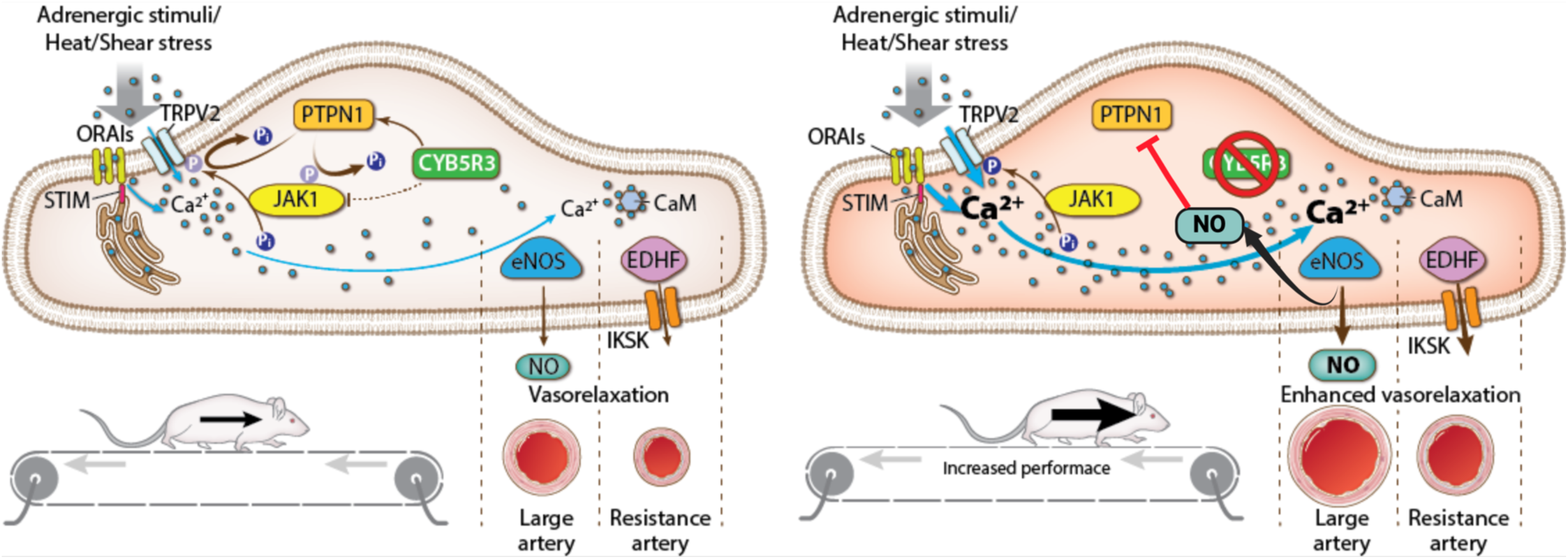
Figure 6. CYB5R3 serves as a redox gatekeeper linking Ca²⁺ influx, phosphatase activity, and NO signaling in the endothelium. **Left:** Under normal conditions, CYB5R3 maintains endothelial redox balance by preserving the catalytic activity of the tyrosine phosphatase PTPN1. Active PTPN1 suppresses TRPV2 activity and limits Ca^2+^ entry through both store-operated (SOCE) and TRPV2-mediated pathways, thereby preserving normal nitric oxide (NO) signaling and vascular tone. **Right:** In CYB5R3-deficient endothelial cells, increased NO-derived oxidants (e.g., S-nitrosothiols, peroxynitrite) inactivate PTPN1, resulting in uncontrolled TRPV2 phosphorylation and increased Ca²⁺ influx. This redox-dependent dysregulation enhances endothelial Ca^2+^ and NO signaling, resulting in vasodilation and improved exercise performance *in vivo*. The schematic illustrates the proposed CYB5R3–PTPN1–TRPV2 axis as a redox-sensitive regulator of Ca^2+^ homeostasis and vascular function, with its imbalance potentially contributing to endothelial dysfunction during aging or in cardiovascular disease.

### Sources of Funding

This work was supported by the National Institutes of Health (NIH) grants: NIH/ National Heart, Lung, and Blood Institute R35HL161177 (A.C.S.), R35HL150778 (M.T.), and 2T32HL110849 – 11A1 (M.K.).

### Disclosures

A.C.S. discloses a financial interest in Creegh Pharmaceuticals Inc. M.T. is a scientific advisor for Seeker Biologics (Cambridge, MA), Eldec Pharmaceuticals (Durham, NC), and Vivreon Biosciences (San Diego, CA).

### Author Contributions

M.K. conceived the study hypothesis, designed and performed the majority of experiments, analyzed data and wrote the manuscript. S.Y. performed the RNAseq experimental design, sample preparation, and data processing. S.N.T. and S.A.H. performed the *ex vivo* wire myography, exercise studies, and maintained the animal colony. R.H. and S.S.T. performed and analyzed molecular assays. O.R. performed the electrophysiology measurements and analyzed the data. K.C.W. designed experiments, conceived the study hypothesis, and wrote the manuscript. A.C.S. and M.T. conceived the study hypothesis, supervised the study, and wrote the manuscript. All authors reviewed and edited the manuscript.

**Supplementary figure 1:**
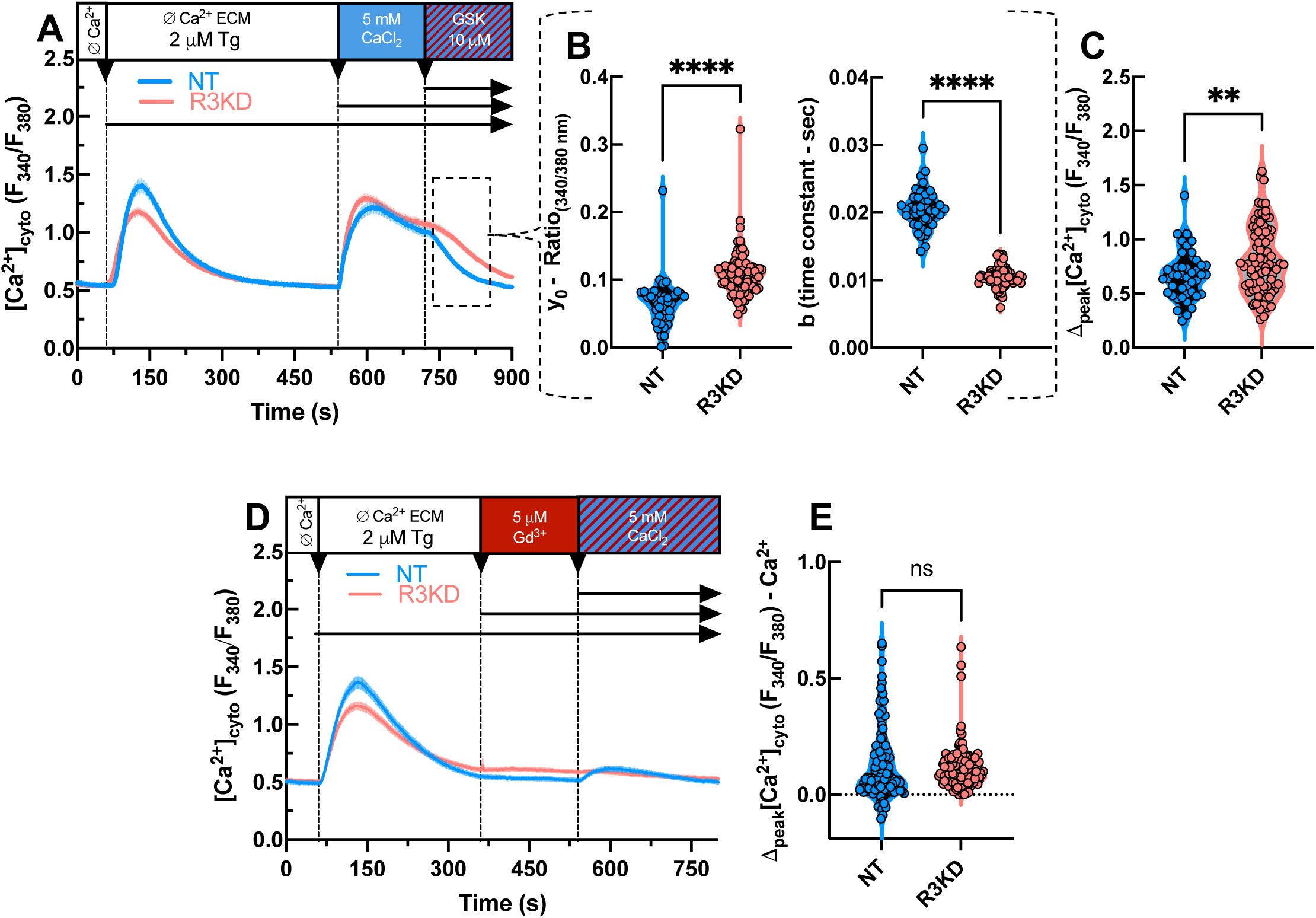
(A) Average time courses showing the changes in [Ca^2+^]_cyto_ (F_340_/F_380_ Ratio) during SOCE protocol in NT (n=43) and R3KD (n=89) HAECs, where (10 μM) GSK-7975A was used to inhibit SOCE. (B) Exponential decay fits (y0, b) on the time courses after GSK-7975A addition. (C) Peak [Ca^2+^]_cyto_ responses after Ca^2+^ re-addition. D) Mean time traces of the changes in the [Ca^2+^]_cyto_ in NT and R3KD HAECs during SOCE protocol, where SOCE was pre-blocked by Gd^3+^ before Ca^2+^ re-addition. E) Delta peak [Ca^2+^]_cyto_ (F_340_/F_380_ Ratio) changes after Ca^2+^ re-addition. Statistics: Welch’s unpaired two-tailed t-test; NT vs R3KD, (B) ****p<0.0001, (C) **p=0.0036, (E) ns p=0.3111.

**Supplementary Figure 2.**
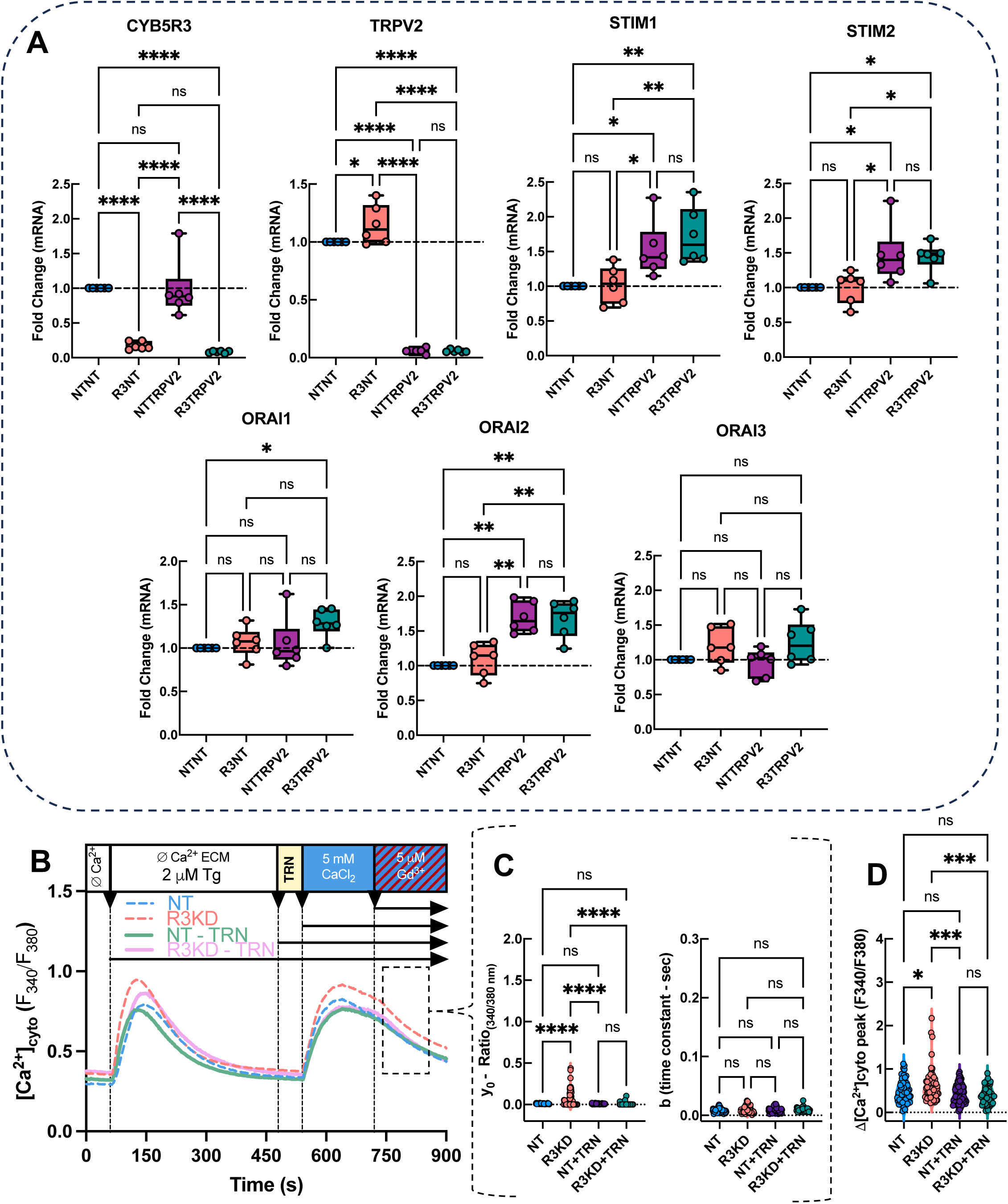
(A) Relative changes in the mRNA of CYB5R3, TRPV2, STIM1 and STIM2, and ORAI1, ORAI2 and ORAI3 in NT&NT, R3KD&NT, NT&TRPV2 and R3&TRPV2KD double siRNA treated cells. B) Time traces (mean ± S.E.M.) showing the changes in [Ca_2+_]_cyto_ (F_340_/F_380_ Ratio) during SOCE protocol in double NT and R3KD HAECs without and with Tranilast (TRN) treatment. HAECs. C) Exponential decay fits showing the y0 and b (time constant) after Gd^3+^-induced inhibition of SOCE (calculated from the first 2 minutes of the inhibition). **Statistics:** (C) One-way ANOVA with Šídák’s multiple comparisons test; (y0) NT vs R3KD, ****p<0.0001; R3KD vs NT+TRN, ****p<0.0001; R3KD vs R3KD+TRN, ****p<0.0001; all other comparisons ns, (b) no significant differences among groups (all p>0.65) ns, (D) NT vs R3KD, *p=0.025; R3KD vs NT+TRN, ***p=0.0003; R3KD vs R3KD+TRN, ***p=0.0002; all other comparisons ns.

**Supplementary Figure 3:**
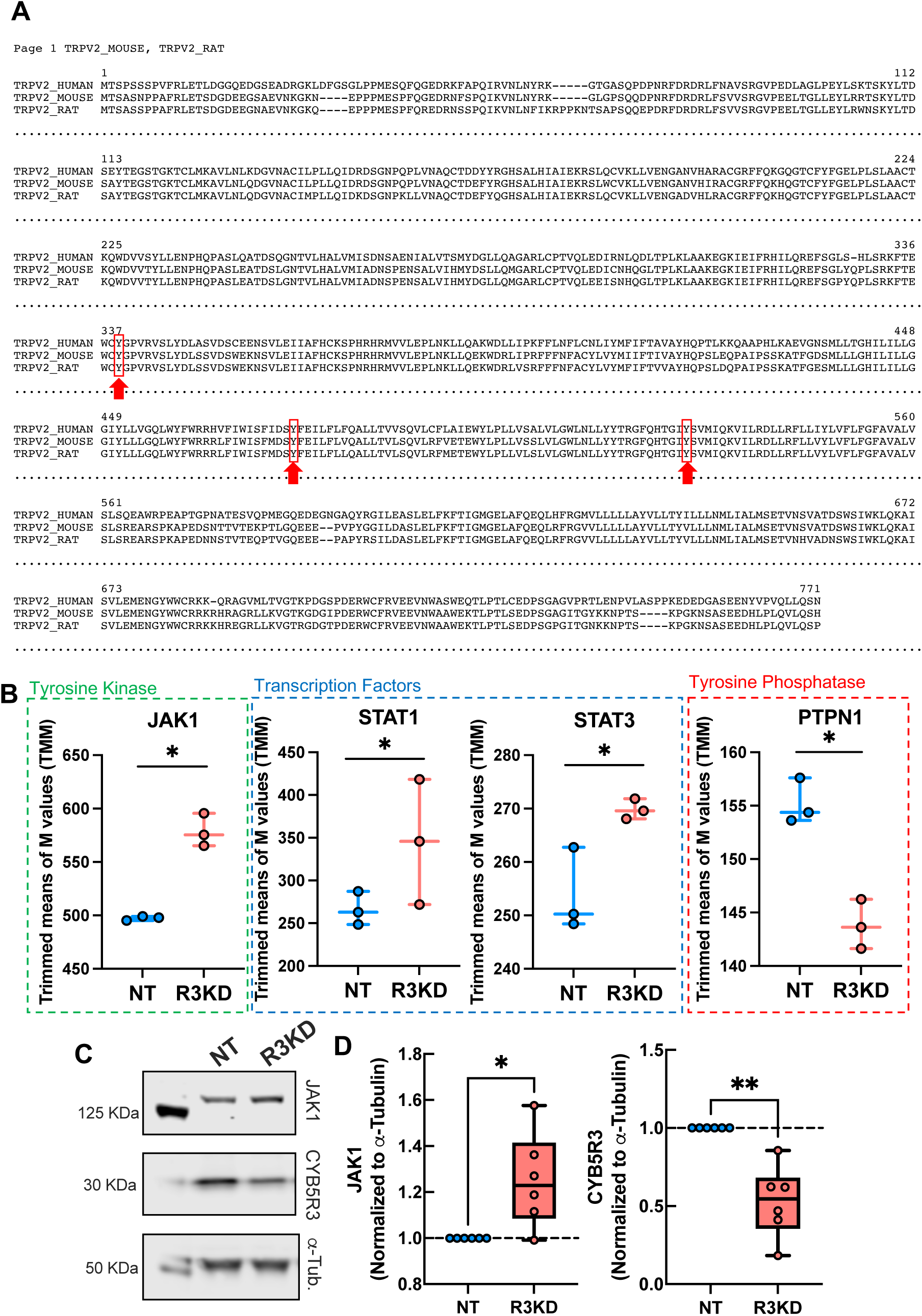
A) Aligned amino acid sequences of human, mouse, and rat TRPV2 orthologs. Preserved tyrosine residues are highlighted with red arrows and boxes. (B) CYB5R3-induced changes in the JAK1, STAT1, STAT3, and PTPN1 in the RNAseq dataset represented as trimmed means of M values. (C) Representative immunoblots showing total JAK1 and CYB5R3 protein expression in NT and CYB5R3K HAEC whole cell lysates. (D) ɑ-Tubulin normalized quantification of JAK1 and CYB5R3 protein levels. **Statistics:** (B) Differential expression was analyzed using DESeq2 (Wald test with Benjamini–Hochberg FDR correction, FDR<0.05) *p<0.001. (D) JAK1: Paired two-tailed t-test, t(5)=3.010, *p=0.0298. CYB5R3: Paired two-tailed t-test, t(5)=5.070, **p=0.0039.

